# Double triage to identify poorly annotated genes in maize: The missing link in community curation

**DOI:** 10.1101/654848

**Authors:** Marcela K. Tello-Ruiz, Cristina F. Marco, Fei-Man Hsu, Rajdeep S. Khangura, Pengfei Qiao, Sirjan Sapkota, Michelle C. Stitzer, Rachael Wasikowski, Hao Wu, Junpeng Zhan, Kapeel Chougule, Lindsay C. Barone, Cornel Ghiban, Demitri Muna, Andrew C. Olson, Liya C. Wang, Doreen C. Ware, David A. Micklos

## Abstract

The sophistication of gene prediction algorithms and the abundance of RNA-based evidence for the maize genome may suggest that manual curation of gene models is no longer necessary. However, quality metrics generated by the MAKER-P gene annotation pipeline identified 17,225 of 130,330 (13%) protein-coding transcripts in the B73 Reference Genome V4 gene set with models of low concordance to available biological evidence. Working with eight graduate students, we used the Apollo annotation editor to curate 86 transcript models flagged by quality metrics and a complimentary method using the Gramene gene tree visualizer. All of the triaged models had significant errors – including missing or extra exons, non-canonical splice sites, and incorrect UTRs. A correct transcript model existed for about 60% of genes (or transcripts) flagged by quality metrics; we attribute this to the convention of elevating the transcript with the longest coding sequence (CDS) to the canonical, or first, position. The remaining 40% of flagged genes resulted in novel annotations and represent a manual curation space of about 10% of the maize genome (~4,000 protein-coding genes). MAKER-P metrics have a specificity of 100%, and a sensitivity of 85%; the gene tree visualizer has a specificity of 100%. Together with the Apollo graphical editor, our double triage provides an infrastructure to support the community curation of eukaryotic genomes by scientists, students, and potentially even citizen scientists.

## Introduction

Maize is the most important cereal crop, with worldwide production nearly equal to wheat and rice tonnage combined [1]. Arguably, only the human genome has received greater scientific scrutiny. The maize genome sequence was published in 2009 [2] and was the last and largest genome generated by the same, laborious clone-by-clone method as the human genome.

Improvements in technology have obviated the requirement of bacterial cloning and decreased DNA sequencing costs 50,000-fold since the initial publication of the maize genome [3,4]. It now costs about $10,000 to generate 50-fold coverage of an average eukaryotic genome, and an additional $20,000 to assemble the millions of individual sequence reads into scaffolds that represent individual chromosomes. However, possession of an assembled genome sequence is only the beginning to understanding an organism’s biology. Genome annotation, and/or curation, adds layers of meaning to the bare sequence of As, Ts, Cs, and Gs. Structural annotation identifies the chromosomal location of a protein-coding gene and creates one or more transcript models of the arrangement of the coding and noncoding information within it. Functional annotation describes elements that control gene transcription, the biological role of the encoded protein, and domains with specific biological activities. This article focuses on protein-coding genes, but genomes also contain transfer RNA genes, transposons, and short-and long-noncoding RNAs.

Protein-coding gene prediction relies of two types of evidence. Mathematical evidence is developed *ab initio* (from the beginning), directly from the assembled genome sequence. Computer algorithms – such as Genefinder, FGenesH, Augustus, and GeneMark –search for patterns in DNA sequence that define a gene, including a start codon, amino acid codons, intron/exon boundaries, and a stop codon. Pattern-based programs typically are trained on a set of representative known genes to develop a hidden Markov model (HMM), which identifies organismal biases for these gene features that are “hidden” in DNA sequence. Biological evidence is provided by experiments that provide mRNA and, to a lesser extent, protein sequences. Homology-based programs look for similarities between the genome sequence and independent RNA and protein evidence from the organism under study and from related organisms. Modern gene annotation programs, such as MAKER-P used for the reference maize genome, employ an iterative process to combine both mathematical and biological evidence to produce increasingly accurate gene models.

Manual annotation, or curation, involves a person evaluating one gene at a time, adding information and making corrections. Annotation jamborees have provided intensive but sporadic annotation efforts. Notably, the *Drosophila melanogaster* genome underwent an early round of annotation by a jamboree of volunteers; community involvement was supported by Apollo, a desktop graphical annotation system [5,6]. Ongoing annotation efforts have focused on humans and model organisms, including GENCODE/Human and Vertebrate Analysis and Annotation (HAVANA) [7]. Organism-specific databases, such as FlyBase (*Drosophila*) [8], WormBase (*Caenorhabditis elegans*) [9], and the Arabidopsis Information Resource (TAIR) [10] – rely primarily on professional curators who focus on functional annotations that add information to the underlying gene model. In contrast, structural curation improves the underlying gene model using additional evidence. Curation of gene models relies mainly on direct input from community members, who discover discrepancies in genes of interest. However, funding for even prominent curation efforts, such as TAIR, is problematic [11], and 62% of biological databases are “dead” in within 18 years [12].

More than a decade after the initial annotation of *Drosophila melanogaster*, all protein-coding genes, long non-coding RNAs, and pseudogenes were manually annotated by FlyBase curators using a Gbrowse genome viewer [13,14]. However, in other organisms, it is often difficult to determine the percentage of gene models that have actually been reviewed by human curators. Since the publication of the *Caenorhabditis elegans* genome two decades ago, curators have set a “Last_reviewed” field for the structures of about 14,000 of 20,000 coding sequences. Although many of the remaining structures may have been looked at by a curator, there is no definitive record of this (Personal communication with G. Williams G, Wormbase, 21 March, 2019). The maize genome was published over a decade ago and has undergone four revisions. Community members used the yrGATE annotation system [15] to curate 231 genes of the B73 RefGen_V2 maize genome [16]. These were the only direct structural improvements hosted on Maize GDB; since that time there has been no organized effort to manually curate maize gene structures (Personal communication with C. Andorf, MaizeGDB, 7 March, 2019). If this is the situation for maize, imagine the status of “orphan genomes” with small research communities. The skeleton in the closet of genome science is that the majority of gene models in the vast majority of sequenced genomes, have not been looked at by any human being – let alone a trained curator. There are good reasons for this.

1. *The volume of genome sequence is overwhelming.* GenBank contains over 500 different eukaryotic genomes that have undergone automated annotation at the National Center for Biotechnology Information (NCBI), with about two new and re-annotated genomes added per week [17]. One of our groups has sequenced 27 maize inbred lines in under a year.
2. *The volume of biological evidence is overwhelming*. The Sequence Read Archive (SRA), the authoritative database for high-throughput data currently has 25.5 quadrillion nucleotides of sequence information and is doubling every 6-8 months [18]. The MaizeCODE Project in which we are involved is developing more than 100 DNA and RNA-seq datasets across five tissues for four maize inbred lines [19]. The scope of curation increases dramatically when one considers that each human gene has an average of four alternative transcripts [7,20], and number for maize still needs to be determined. Each RNA-seq experiment from a different tissue or developmental timepoint potentially adds new isoforms. This presents a moving target of increasing numbers of alternatively spliced transcripts.
3. *Automated gene annotation seems good enough.* Retrospective studies in the human genome have shown that HMMs can correctly identify about 85% of individual exons and every exon in about 58% of protein-coding genes [21]. An analysis in bread wheat revealed FGenesH as the best gene finder, predicting more than 75% of all the genes correctly [22]. Automated annotation continues to improve with the increasing availability of RNA-seq and long-read RNA evidence from single molecule sequencing platforms produced by Pacific Biosciences and Oxford Nanopore [23]. However, the rapid accumulation of automated genome annotations creates additional problems, as errors in draft sequences are propagated to orthologous genes in other species [24].
4. *There has been little guidance on where to focus effort on structural annotations.* Given the fact that most gene models are correct or nearly correct, there is little potential reward in inspecting random genes. To date, there have been no recommendations on how to identify genes in need of manual curation.

Community annotation by students and non-expert researchers is held out as a means to curate the growing number of sequenced eukaryotic genomes, most of which lack dedicated funding or database resources. Manual curation provides an ideal way to give students an intuitive understanding of gene structure and function, while providing researchers with high-quality genome data [25]. The Genomic Education Partnership (GEP) involved hundreds of undergraduate students in manually annotating genes on the *Drosophila* Muller F elements [26]. In another project, undergraduate and graduate students worked with experienced curators to annotate 530 genes in *Diaphorina citri* Kuwayama. This non-model insect is the vector of *citrus greening* disease that threatens agriculture worldwide [27].

Our needs assessment showed that maize biologists would like to ensure the accuracy of models for genes they work with and are willing to help out with manual curation. Of 112 PIs, postdocs, and students we surveyed at the 2017 Maize Genetics Conference, 90% said that manual annotations of maize gene families would be useful to their research; 60% would participate in annotating genes of which they had expert knowledge; and 41% would annotate genes as a class project (see S1 Appendix).

Given the high accuracy of most automated annotation and the fact that maize genome is supported by abundant RNA-seq and long-read RNA evidence, we wondered: How can we focus on genes and transcripts most in need of manual curation? As a corollary, how can we support maize researchers and undergraduate faculty in a community annotation effort? We addressed these questions by developing methods to triage maize gene models to identify suspect annotations using MAKER-P quality metrics and the Gramene gene tree visualizer. Then we enlisted the help of young biologists to edit the triaged genes with a web-enabled version of Apollo.

## Results

### Single Triage of 47 Genes from Five Maize Gene Families

We analyzed gene models from the reference sequence of maize B73 (B73 RefGen_V4) [28]. This assembly was annotated with MAKER-P [29], which generates quality metrics that assess how well a transcript model is supported by available biological evidence. We used two of these metrics to identify low-quality gene models: Annotation Edit Distance (AED) and Quality Index 2 (QI2). AED values range between 0 and 1, with 0 denoting perfect concordance with the available evidence and 1 denoting absence of supporting evidence. One of nine Quality Indices generated by MAKER-P, QI2 is the fraction of splice sites confirmed by alignments to RNA evidence. Gene models without introns (therefore no splice sites) are given a QI2 value of 0 [30]. Some gene models, particularly those generated by *ab initio* methods, can show low metrics due to a lack of supporting evidence. To ensure the availability of evidence to use in manual curation, we flagged genes with AED scores less than 0.5 (AED < 0.5) and QI2 values between 0.33 and 0.75 (QI2 0.33-0.75). Applying these quality metrics identified 17,225 of 130,330 (13%) protein-coding transcripts in the B73 RefGen_V4 with low concordance to available biological evidence.

We tested the utility of this triage in a mini-annotation jamboree held in December 2017 at Cold Spring Harbor Laboratory (CSHL). We reasoned that participants would be more engaged by working with genes related to their own research or with obvious biological significance. Therefore, we focused on five well-known gene families: PIN-formed (*PIN*), Gretchenhagen-3 (*GH3*), ATP-binding cassette (*ABC*), cycloid and teosinte branched (*TCP*) and origin recognition complex (*ORC*). During the two-day event, nine graduate students and one postdoctoral fellow examined 40 genes having four or fewer transcripts. This resulted in the curation of 57 transcripts from these genes families, including 11 transcripts flagged by quality metrics and two unflagged transcripts. The transcripts were edited in Apollo, the graphical annotation editor developed to curate the *Drosophila* genome. We used the web-enabled version, which is significantly easier to use and readily supports community annotation [31].

All 11 of the flagged models required curation (Table 1). Six (55%) were missing one or more exons, while two (18%) had an extra exon. Nine (82%) had exons with incorrect lengths, including two (18%) with non-canonical splice sites. Fig 1 provides an example of an exon curation in the PIN family. We were able to extend untranslated regions (UTRs) in three (27%) of the transcripts. Six (55%) of the curated models matched another transcript model for the same gene. Expressed another way, 46% of curations were “novel” (see S2 Table).

**Table 1.**
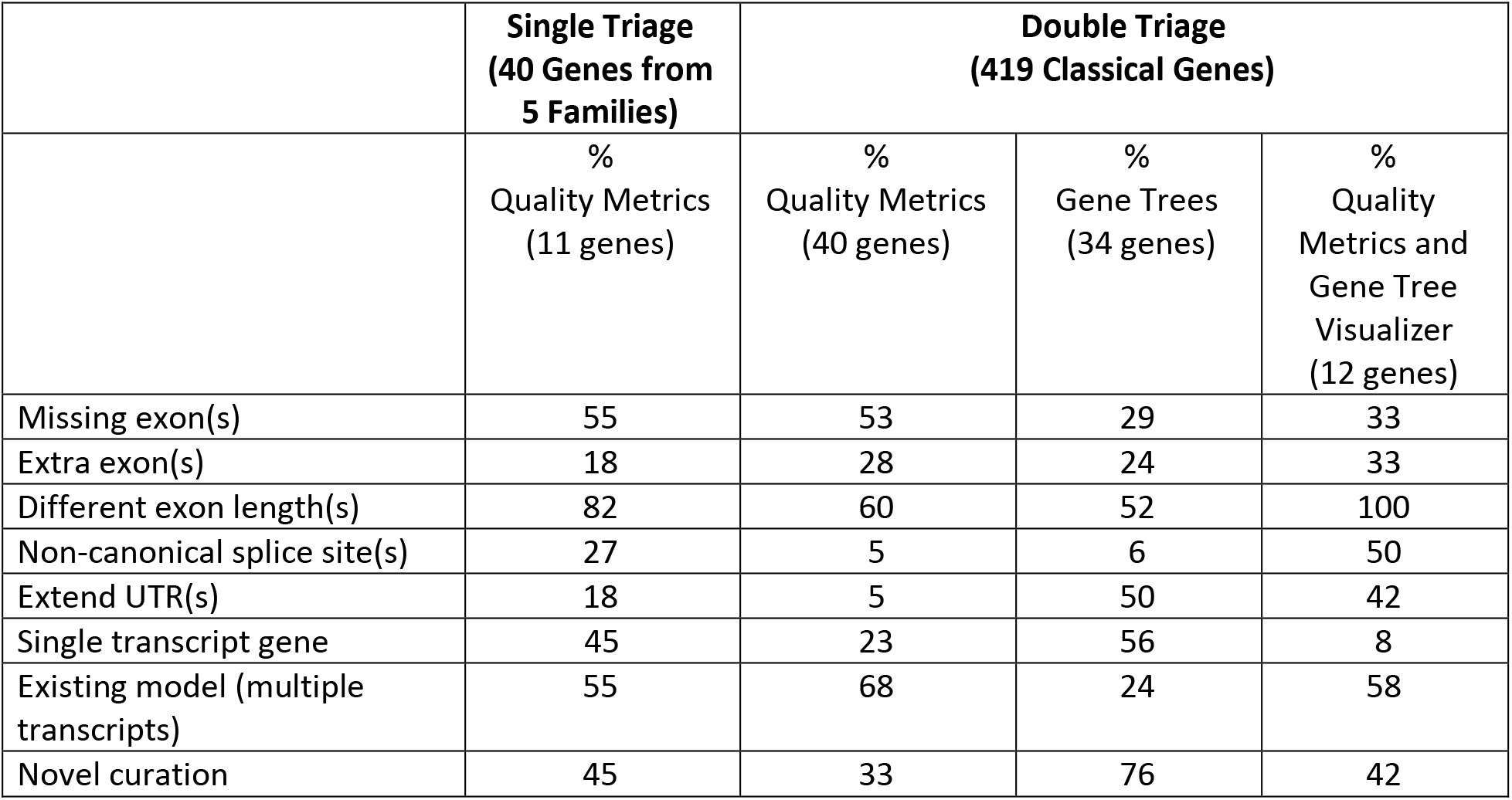
Annotation errors found by single and double triage of maize genes. Errors found in 11 genes flagged by quality metrics in five maize gene families (25 transcripts). Errors found in 40 genes flagged by quality metrics, 34 genes flagged by gene trees and 12 genes flagged by both methods in 419 maize classical genes (2,127 transcripts).

|  | Single Triage<br>(40 Genes from<br>5 Families) | Double Triage<br>(419 Classical Genes) |  |  |
| --- | --- | --- | --- | --- |
|  | %<br>Quality Metrics<br>(11 genes) | %<br>Quality Metrics<br>(40 genes) | %<br>Gene Trees<br>(34 genes) | %<br>Quality<br>Metrics and<br>Gene Tree<br>Visualizer<br>(12 genes) |
| Missing exon(s) | 55 | 53 | 29 | 33 |
| Extra exon(s) | 18 | 28 | 24 | 33 |
| Different exon length(s) | 82 | 60 | 52 | 100 |
| Non-canonical splice site(s) | 27 | 5 | 6 | 50 |
| Extend UTR(s) | 18 | 5 | 50 | 42 |
| Single transcript gene | 45 | 23 | 56 | 8 |
| Existing model (multiple transcripts) | 55 | 68 | 24 | 58 |
| Novel curation | 45 | 33 | 76 | 42 |

**Figure 1.**
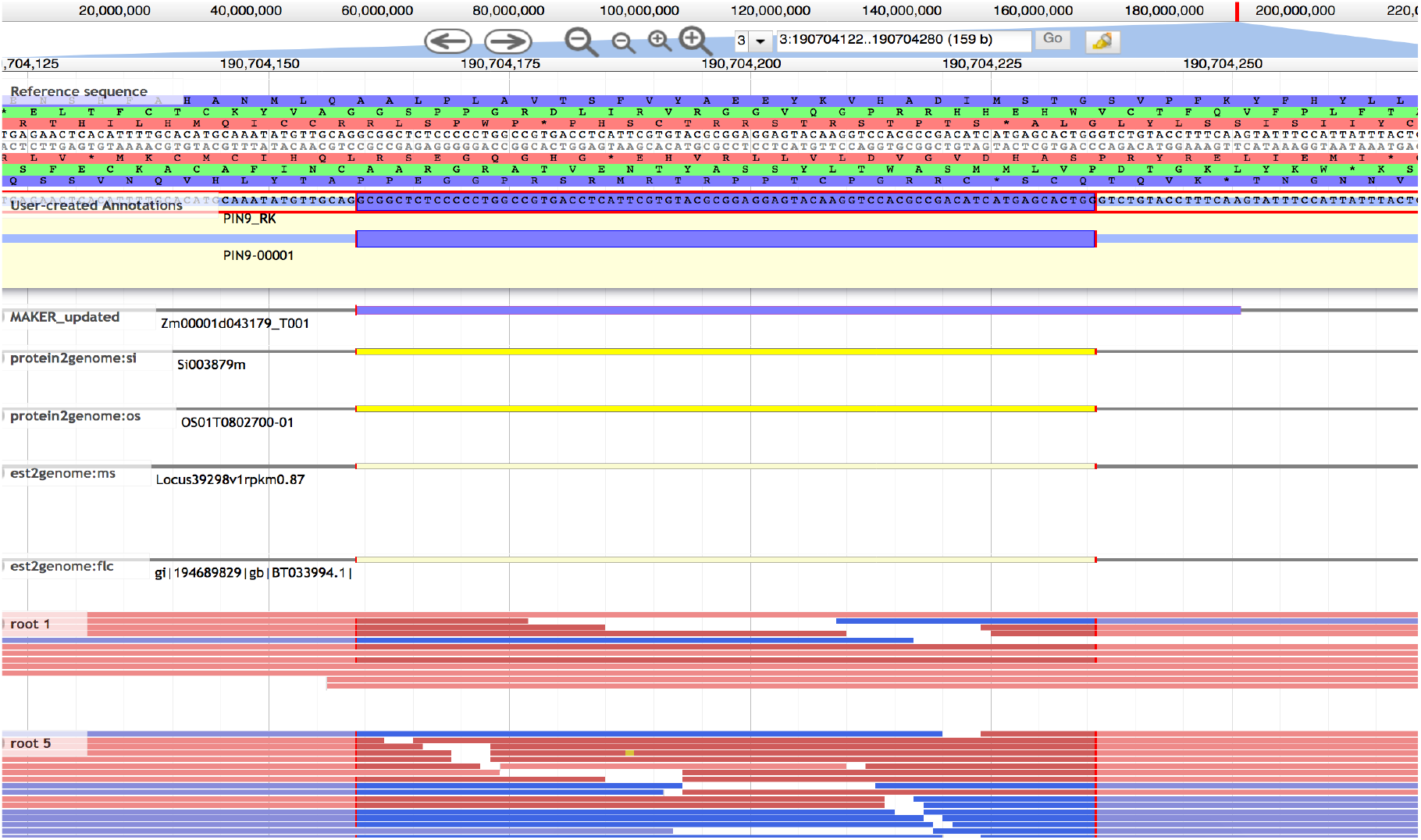
Curation of exon 3 of PIN9 (Zm00001d043179). Exons of incorrect length were the most common error detected by both triage methods. The Apollo editing window shows a “User-created annotation” at top followed by the longer, incorrect B73 RefGen V4 model (“MAKER_updated”). The shortened exon was supported by aligned evidence: protein sequences from sorghum and rice, assembled long Iso-Seq reads combined from six tissues, and RNA-seq from roots, among other tissues.

### Double Triage of 419 Classical Maize Genes

We extended our analysis to a set of “classical” maize genes, which represent well-studied genetic loci that have been cloned [32]. We used an updated list at MaizeGDB [33]. About one quarter of classical genes were first identified by a visible mutant phenotype – and include many markers used to make genetic maps before the availability of molecular markers. We analyzed the canonical (longest protein-coding) transcript of 419 classical genes having 2,127 transcripts and 277 distinct families. Jamboree group members inspected the classical gene models with the gene tree visualizer at Gramene, a comparative database with 58 plant genomes (http://www.gramene.org) [34]. This tool displays a phylogenetic tree and alignments between the translated protein sequences of one classical gene and its homologs across species. Discrepancies in alignments are shown as insertions or deletions. Each week, 10-20 classical genes were triaged independently by three group members using the gene tree visualizer. Fig 2 provides an example of a classical gene flagged using the gene tree visualizer.

**Figure 2.**
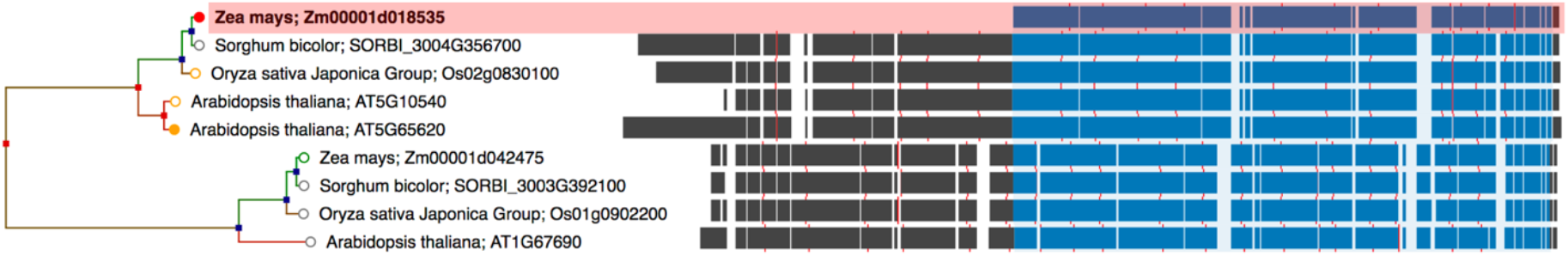
Triage of BRICK1 (Zm00001d042475) with the Gramene gene tree visualizer. Comparison to closest plant orthologs and maize paralogs revealed that the B73 RefGen V4 model was missing the entire 5’ end.

We then curated 34 classical genes flagged by the gene tree method, along with 40 genes flagged by MAKER-P quality metrics, and 12 genes flagged by both methods. This constituted a double triage. Over a 12-month period, we performed multiple cycles of internal review of the curated gene models to troubleshoot methods, identify errors, and suggest improvements to annotations. Curators presented their models for peer review during periodic video conferences. We found errors in 86 (21%) of the classical gene set (see S3 Table). Table 1 compares the errors identified by each triage. Fig 3 shows the different but overlapping sets of genes identified for manual curation by quality metrics and gene trees visualization.

**Figure 3.**
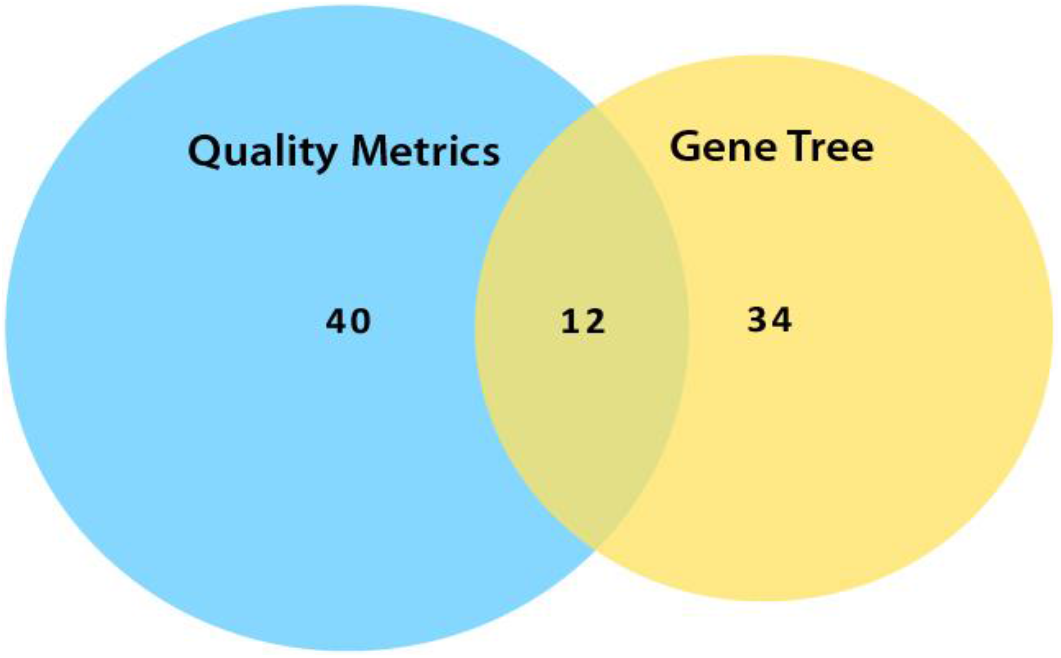
Double triage of classical maize genes. Number of classical gene transcripts with annotation errors flagged by MAKER-P quality metrics and Gramene gene tree visualizer.

### Curation of an exceptional gene family

Five of the 12 maize genes in the acidic invertase gene family were included in the classic gene set we curated. Two members were flagged by MAKER-P quality metrics and annotated in Apollo (INVVR2 and INVCW3). Prompted by a recent report of potential annotation errors in members of this family [35], we undertook an in-depth evaluation of all 12 family members (see S4 Table). Alignment with available evidence confirmed the presence of a 9-nucleotide mini-exon encoding a tripeptide that had been included in B73 RefGen_V3 models for six family members. This conserved DPN peptide is predicted to be the active site of a β-fructosidase or invertase [36–39]. We discovered this mini-exon in four additional family members, which had not been previously reported. We also identified a novel 19-nt exon in INVCW4, a cell wall invertase (Fig 4).

**Figure 4.**
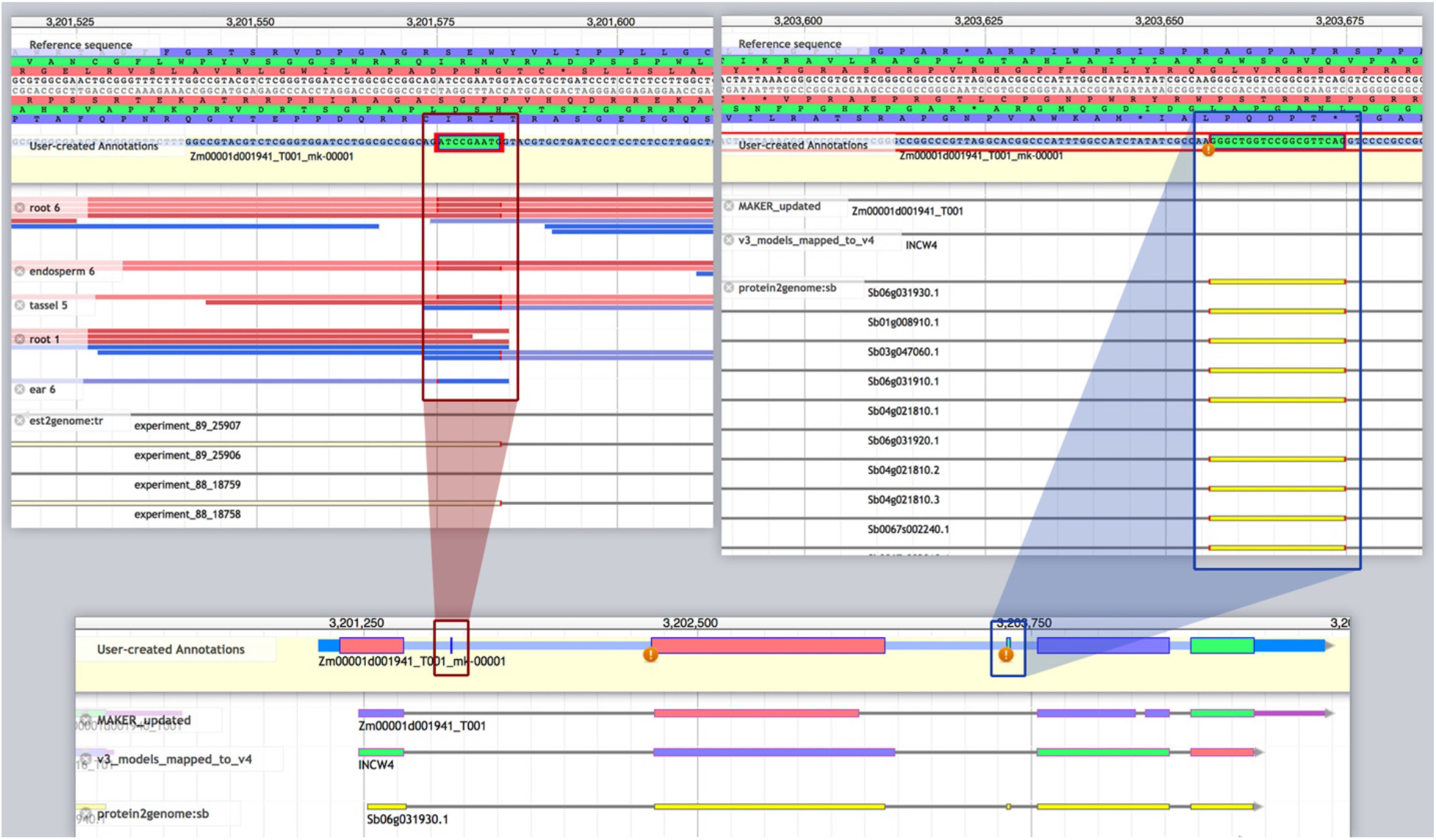
Curation of two mini-exons in INVCW2 (Zm00001d043179). The Apollo editing window shows a “User-created annotation” at top, followed by incorrect B73 RefGen V4 model (“MAKER_updated”) and “v3 model mapped to v4.” A conserved 9-nucleotide exon (red circle) and a novel 19-nucleotide exon (blue circle) were supported by protein sequences from sorghum and rice, assembled EST transcripts from ultra-deep sequencing, long Iso-Seq reads combined from six tissues, and RNA-seq from root and other tissues.

## Discussion

We have demonstrated that there is significant room for improving the annotations of even well documented protein-coding genes within a well-studied genome, such as maize. Our studies show that MAKER-P quality metrics and the Gramene gene tree visualizer offer effective and complementary triages to identify poor-quality gene models. All of the genes flagged by quality metrics and gene trees required curation; a specificity of 100% for both methods. Sensitivity was more difficult to assess with our data set, but quality metrics had a sensitivity of 85% to detect annotation errors.

Each triage has its strengths and weaknesses. MAKER-P quality metrics can be used to quickly generate list of suspected genes in any genome. The majority of genes flagged by this method had multiple transcripts, and the curated models frequently matched an existing transcript model in the v4 gene set. So, this triage produced a lower percentage of novel curations. The gene tree visualizer takes more time, but provides a wholistic, phylogenetic approach to curation. This method aligns related orthologs and performed well with the classical genes, which tend to be highly conserved across grasses. The majority of genes identified by gene trees had a single transcript and produced novel annotations. Gene trees picked up genes missed by quality metrics triage, and thus provides a complement to the automated method.

Our initial triage of the B73 RefGen_V4 gene set with quality metrics flagged 13% of the protein-coding transcripts for potential annotation errors. However, a correct annotation existed for about 60% of the flagged transcripts we edited. The V4 gene annotation used the Ensembl platform [40], which set the isoform with the longest coding sequence (CDS) as the canonical transcript. Our study suggests that a majority of transcripts elevated to canonical status on length alone are incorrect and that the transcript with the lowest AED is the best choice for a canonical model. This explains why in the gene family analysis, in most cases, the transcript with the lowest AED was the correct one. We will use this information to tune MAKER-P for the annotation of maize version 5, and we recommend that quality metrics be considered in selecting the primary transcript in other genomes. In this way curation can provide feedback to make informed updates to automated annotation systems.

We have demonstrated that there is interest in community annotation, and we have provided a method to make this possible. Researchers are willing to commit time to manual curation, because they realize the negative impact of poor models to their research. Improvement of reference sequences will be especially important as we move toward synthetic biology. We believe that our dual triage is the missing link in popularizing community and even citizen science annotation. It focuses on the fraction of gene models that demands attention. Gene triaging generates and maintains interest by ensuring that each curation effort will be rewarded with new contributions to genome science.

## Materials and Methods

We surveyed 119 attendees of the 59^th^ Annual Maize Genetics Conference, March 10-12, 2017. Participants were selected randomly and asked to confidentially complete an online questionnaire on a handheld tablet. The results were tabulated in Survey Monkey and analyzed using IBM SPSS Statistics 23. All survey activities were reviewed and approved by the Cold Spring Harbor Laboratory Institutional Review Board (IRB no. 17-007).

The B73 reference sequence (B73 RefGen_V4) was annotated with MAKER-P version 3.1 [28]. This bioinformatics pipeline integrated *ab initio* gene prediction with publicly available evidence from full-length cDNA [41], *de novo* assembled transcripts from short-read mRNA sequencing (RNA-seq) [42], isoform-sequencing (Iso-Seq) full-length transcripts [43], and proteins from *Sorghum bicolor*, *Oryza sativa*, *Setaria italica*, *Brachypodium distachyon*, and *Arabidopsis thaliana* [34].

Apollo (http://genomearchitect.github.io/) is a genome annotation platform originally developed to support annotation of the *Drosophila* genome. The latest version of Apollo is web-based and built on the popular JBrowse genome browser. Apollo displays as features experimental data (e.g. RNA-seq data, cDNAs, or other imported data sets) as well as predictions from gene annotation pipelines (e.g. transcripts, variant calls, repeat regions, etc.). The features (available in the “Evidence Area” of the interface) can be imported from any of several file formats, including GFF3, BAM, GTF, GVF, GenBank, BED, BigWig, or Chado database. Users can drag one or more features from the “Evidence Area” to the “Editing Area” to synthesize a new or refined annotation of a gene. Most editing is done through intuitive drag or drag-and-drop manipulations. Additional menus manage display parameters. Apollo records a complete editing history of a user-created annotation, and also allows for real-time collaboration on a project. Apollo’s rich set of features offers a scalable and integrated platform that has supported several community annotation efforts.

The Gramene gene tree visualizer provides an interactive interface to inspect protein sequence alignments for a given gene family and identify genes with potential annotation errors. Alignments are shown as the branches of a phylogenetic tree centered on a gene of interest, and a simple click allows the tree to be rearranged around the center on a different gene within the same tree. Three display modes allow the trees to be explored at various levels: 1) In the Alignment Overview, InterPro Scan descriptions are accessed by clicking on color-coded domains, 2) in the Multiple Sequence Alignment view, a slider is dragged to scan the amino sequence and select a standard color schema (such as Clustal, Zappo, or helix propensity), 3) In the Neighborhood Conservation view, 10 flanking genes are displayed on each side of the gene of interest, color-coded by gene family. For this project, we extended the interface to let users flag genes for further curation (http://curate.gramene.org). We set up a python/flask web service and database to store and review the results. Phylogenetic trees available in the viewer were generated via the Ensembl Compara pipeline [43] using amino acid sequences from 52 species in Gramene build 56.

All of our novel annotations are available as a separate track (“curated_apollo_annotations”) in the Gramene browser. Go to http://news.gramene.org/curated_maize_v4_gene_models, and click on the “Genomic coordinates” of a gene of interest. <u>This will pull up a Gramene browser window centered on that gene. Scroll down</u> to scroll down to view the gene models. Our annotations are also available in gff3 format at Track Hub Registry (ftp://ftp.gramene.org/pub/gramene/CURRENT_RELEASE/gff3/zea_mays/apollo_annotations_maize_v4.gff3). A complete list of the 2,127 transcripts used in this analysis can be found in S5 Table).

## Acknowledgments

We thank Michael Campbell for suggesting the MAKER-P parameters we used to flag suspect annotations. We thank Suzi Lewis and Ed Lee for creating Apollo, for evolving it into its current web-enabled form, and for helping us adapt it for use in education outreach. We thank Monica Munoz-Torres for providing an annotation tutorial for the jamboree.

## Author Contributions

Conceived and designed the experiments: DW, DAM, MKT-R, CFM. Performed the experiments: MKT-R, CFM, RSK, MCS, SS, HW, JZ, RW, PQ, F-MH, LCB, DM, CG, ACO, LCW, KC. Analyzed the data: MK-T, CFM, DAM. Wrote the paper: DAM, MKT-R, CFM, DW.

## References

1. Foreign Agricultural Service, United States Department of Agriculture. All grain summary comparison [Internet]. 2019. Available at https://apps.fas.usda.gov/psdonline/circulars/grain.pdf (p. 15)

2. Schnable PS, Ware D, Fulton RS, Stein JC, Wei F, Pasternak S, et al. The B73 maize genome: complexity, diversity, and dynamics. Science. 2009;326: 1112–1115. doi:10.1126/science.1178534.

3. National Human Genome Research Institute. Cost per raw megabase of DNA sequence. 2017. Available at https://www.genome.gov/images/content/costpermb_2017.jpg

4. Barone L, Williams J, Micklos D. Unmet needs for analyzing biological big data: A survey of 704 NSF principal investigators. PLS Comput Biol. 2017;13: e1005755. doi:10.1371/journal.pcbi.1005755

5. Pennisi E. Ideas fly at gene-finding jamboree. Science. 2000;287: 2182–2184. Available at https://www.ncbi.nlm.nih.gov/pubmed/10744542

6. Misra S, Crosby MA, Mungall CJ, Matthews BB, Campbell KS, Hradecky P, et al. Annotation of the Drosophila melanogaster euchromatic genome: a systematic review. Genome Biol. 2002;3: RESEARCH0083. Available at https://www.ncbi.nlm.nih.gov/pubmed/12537572

7. Harrow J, Frankish A, Gonzalez JM, Tapanari E, Diekhans M, Kokocinski F, et al. GENCODE: the reference human genome annotation for The ENCODE Project. Genome Res. 2012;22: 1760–1774. doi:10.1101/gr.135350.111

8. Thurmond J, Goodman JL, Strelets VB, Attrill H, Gramates LS, Marygold SJ, et al. FlyBase 2.0: the next generation. Nucleic Acids Res. 2019;47: D759–D765. doi:10.1093/nar/gky1003

9. Harris TW, Chen N, Cunningham F, Tello-Ruiz M, Antoshechkin I, Bastiani C, et al. WormBase: a multi-species resource for nematode biology and genomics. Nucleic Acids Res. 2004;32: D411–7. doi:10.1093/nar/gkh066

10. Berardini TZ, Reiser L, Li D, Mezheritsky Y, Muller R, Strait E, et al. The Arabidopsis information resource: Making and mining the “gold standard” annotated reference plant genome. Genesis. 2015;53: 474–485. doi:10.1002/dvg.22877

11. Reiser L, Berardini TZ, Li D, Muller R, Strait EM, Li Q, et al. Sustainable funding for biocuration: The Arabidopsis Information Resource (TAIR) as a case study of a subscription-based funding model. Database. 2016. 2016. doi:10.1093/database/baw018

12. Attwood TK, Agit B, Ellis LBM. Longevity of Biological Databases. EMBnet.journal. 2015;21: 803. doi:10.14806/ej.21.0.803.

13. Crosby MA, Gramates LS, Dos Santos G, Matthews BB, St Pierre SE, Zhou P, et al. Gene Model Annotations for Drosophila melanogaster: The Rule-Benders. G3. 2015;5: 1737–1749. doi:10.1534/g3.115.018937

14. Matthews BB, Dos Santos G, Crosby MA, Emmert DB, St Pierre SE, Gramates LS, et al. Gene Model Annotations for Drosophila melanogaster: Impact of High-Throughput Data. G3. 2015;5: 1721–1736. doi:10.1534/g3.115.018929

15. Wilkerson MD, Schlueter SD, Brendel V. yrGATE: a web-based gene-structure annotation tool for the identification and dissemination of eukaryotic genes. Genome Biol. 2006;7: R58. doi:10.1186/gb-2006-7-7-r58

16. Available at http://www.plantgdb.org/ZmGDB/DisplayProjects.php

17. Eukaryotic Genome Annotation at NCBI. Available at [Internet]. Available at https://www.ncbi.nlm.nih.gov/genome/annotation_euk/

18. Sequence Read Archive. National Center for Biotechnology Information. Available at. https://trace.ncbi.nlm.nih.gov/Traces/sra/sra.cgi?view=announcement.

19. Available at https://www.nsf.gov/awardsearch/showAward?AWD_ID=1445025

20. GENCODE. Statistics about the current GENCODE Release (version 29). Available at https://www.gencodegenes.org/human/stats.html.

21. Kulp D, Haussler D, Reese MG, Eeckman FH. A generalized hidden Markov model for the recognition of human genes in DNA. Proc Int Conf Intell Syst Mol Biol. 1996;4: 134–142. Available at https://www.ncbi.nlm.nih.gov/pubmed/8877513

22. Nasiri J, Naghavi M, Rad SN, Yolmeh T, Shirazi M, Naderi R, et al. Gene identification programs in bread wheat: a comparison study. Nucleosides Nucleotides Nucleic Acids. 2013;32: 529–554. doi:10.1080/15257770.2013.832773

23. Weirather JL, de Cesare M, Wang Y, Piazza P. Comprehensive comparison of Pacific Biosciences and Oxford Nanopore Technologies and their applications to transcriptome analysis. ncbi.nlm.nih.gov; 2017. Available at https://www.ncbi.nlm.nih.gov/pmc/articles/PMC5553090.2/

24. Salzberg SL. Next-generation genome annotation: we still struggle to get it right. Genome Biology. 2019;20 (92). doi:10.1186/s13059-019-1715-2.

25. Hosmani PS, Shippy T, Miller S, Benoit JB, Munoz-Torres M et al. A quick guide for student-driven community genome annotation. PLoS Comput. Biol. 2019; 15(4):e1006682. doi:10.1371/journal.pcbi.1006682

26. Leung W, Shaffer CD, Reed LK, Smith ST, Barshop W, Dirkes W, et al. Drosophila muller f elements maintain a distinct set of genomic properties over 40 million years of evolution. G3. 2015;4;5(5):719–40. doi:10.1534/g3.114.015966.

27. Saha S, Hosmani PS, Villalobos-Ayala K, Miller S, Shippy T, Flores M et al. Improved annotation of the insect vector of citrus greening disease: biocuration by a diverse genomics community. Database. 2019. 2019. doi:10.1093/database/baz035.

28. Jiao Y, Peluso P, Shi J, Liang T, Stitzer MC, Wang B, et al. Improved maize reference genome with single-molecule technologies. Nature. 2017;546: 524–527. doi:10.1038/nature22971

29. Campbell MS, Holt C, Moore B, Yandell M. Genome Annotation and Curation Using MAKER and MAKER-P. Curr Protoc Bioinformatics. 2014;48: 4.11.1–39. doi:10.1002/0471250953.bi0411s48

30. Eilbeck K, Moore B, Holt C, Yandell M. Quantitative measures for the management and comparison of annotated genomes. BMC Bioinformatics. 2009;10: 67. doi:10.1186/1471-2105-10-67

31. Dunn NA, Unni DR, Diesh C, Munoz-Torres M, Harris NL, Yao E, et al. Apollo: Democratizing genome annotation. PLoS Comput Biol. 2019;15: e1006790. doi:10.1371/journal.pcbi.1006790

32. Schnable JC, Freeling M. Genes identified by visible mutant phenotypes show increased bias toward one of two subgenomes of maize. PLoS One. 2011;6: e17855. doi:10.1371/journal.pone.0017855

33. Available at https://www.maizegdb.org/associated_genes?type=classical&style=table

34. Tello-Ruiz MK, Naithani S, Stein JC, Gupta P, Campbell M, Olson A, et al. Gramene 2018: unifying comparative genomics and pathway resources for plant research. Nucleic Acids Res. 2018;46: D1181–D1189. doi:10.1093/nar/gkx1111

35. Juárez-Colunga S, López-González C, Morales-Elías NC, Massange-Sánchez JA, Trachsel S, Tiessen A. Genome-wide analysis of the invertase gene family from maize. Plant Mol Biol. 2018;97: 385–406. doi:10.1007/s11103-018-0746-5

36. Sturm A. Invertases. Primary structures, functions, and roles in plant development and sucrose partitioning. Plant Physiol. 1999;121: 1–8. Available at https://www.ncbi.nlm.nih.gov/pubmed/10482654

37. Verhaest M, Lammens W, Le Roy K, De Coninck B, De Ranter CJ, Van Laere A, et al. X-ray diffraction structure of a cell-wall invertase from Arabidopsis thaliana. Acta Crystallogr D Biol Crystallogr. 2006;62: 1555–1563. doi:10.1107/S0907444906044489

38. Yao Y, Geng M-T, Wu X-H, Liu J, Li R-M, Hu X-W, et al. Genome-wide identification, 3D modeling, expression and enzymatic activity analysis of cell wall invertase gene family from cassava (Manihot esculenta Crantz). Int J Mol Sci. Multidisciplinary Digital Publishing Institute; 2014;15: 7313–7331. Available at https://www.mdpi.com/1422-0067/15/5/7313/htm

39. Yao Y, Geng M-T, Wu X-H, Liu J, Li R-M, Hu X-W, et al. Genome-Wide Identification, Expression, and Activity Analysis of Alkaline/Neutral Invertase Gene Family from Cassava (Manihot esculenta Crantz). Plant Mol Biol Rep. 2015;33: 304–315. doi:10.1007/s11105-014-0743-z

40. Cunningham F, Achuthan P, Akanni W, Allen J, Amode MR, Armean IM, et al. Ensembl 2019. Nucleic Acids Res. 2019;47: D745–D751. doi:10.1093/nar/gky1113

41. Soderlund C, Descour A, Kudrna D, Bomhoff M, Boyd L, Currie J, et al. Sequencing, mapping, and analysis of 27,455 maize full-length cDNAs. PLoS Genet. 2009;5: e1000740. doi:10.1371/journal.pgen.1000740

42. Law M, Childs KL, Campbell MS, Stein JC, Olson AJ, Holt C, et al. Automated update, revision, and quality control of the maize genome annotations using MAKER-P improves the B73 RefGen_v3 gene models and identifies new genes. Plant Physiol. 2015;167: 25–39. doi:10.1104/pp.114.245027

43. Wang B, Tseng E, Regulski M, Clark TA, Hon T, Jiao Y, et al. Unveiling the complexity of the maize transcriptome by single-molecule long-read sequencing. Nat Commun. 2016;7: 11708. doi:10.1038/ncomms11708

44. Herrero J, Muffato M, Beal K, Fitzgerald S, Gordon L, Pignatelli M, et al. Ensembl comparative genomics resources. Database. 2016;2016. doi:10.1093/database/baw053

