## Supplementary material for "Double triage to identify poorly annotated genes in maize: The missing link in community curation"

### S1: Appendix, Survey Results

#### Table 1: Inbred lines used in survey respondent research

| | Count | % |
|---|---|---|
| B73 | 98 | 87.50% |
| W22 | 64 | 57.14% |
| NC350 | 23 | 20.54% |
| Teosinte inbred (TIL11) | 22 | 19.64% |
| None of the above | 11 | 9.82% |

*n = 112, participants could select more than one option*

#### Table 2: Tissues used in survey respondent research

| | Count | % |
|---|---|---|
| Immature ears | 48 | 43.24% |
| Root tips | 44 | 39.64% |
| Roots | 43 | 38.74% |
| Shoot tips | 40 | 36.04% |
| Mature pollen | 40 | 36.04% |
| Embryo 15 dap | 38 | 34.23% |
| Endosperm 15 dap | 36 | 32.43% |
| FACS-sorted cell types | 22 | 19.82% |
| None of the above | 24 | 21.62% |

*n = 111, participants could select more than one option*

#### Table 3: Data sets important to respondent research

| | Count | % |
|---|---|---|
| RNA-seq | 95 | 84.82% |
| Whole genome sequences | 89 | 79.46% |
| Transcription factor binding sites | 66 | 58.93% |
| Open chromatin marks | 51 | 45.54% |
| DNA methylation sites | 51 | 45.54% |
| Small RNA-seq | 47 | 41.96% |
| Histone modifications | 43 | 38.39% |
| RNA polymerase II binding sites | 32 | 28.57% |
| None of the above | 4 | 3.57% |

*n = 112, participants could select more than one option*

**Table 4: Types of anticipated analyses respondents would do with students**

| | Individual Projects | | Class Projects | |
|---|---|---|---|---|
| | Count | % | Count | % |
| Compare regulatory elements of different inbred lines | 58 | 87.88% | 17 | 25.76% |
| Compare genome structure of different inbred lines | 70 | 87.50% | 21 | 26.25% |
| Compare members of specific gene families | 67 | 85.90% | 27 | 34.62% |
| Compare transposons of different inbred lines | 43 | 82.69% | 14 | 26.92% |

*n = 95, participants could select more than one option.*

**Table 5: Utility of hand-edited maize gene family annotations for research**

| | Count | % |
|---|---|---|
| Not useful | 2 | 1.82% |
| Slightly useful | 9 | 8.18% |
| Useful | 34 | 30.91% |
| Moderately useful | 23 | 20.91% |
| Extremely useful | 42 | 38.18% |

**Table 6: Willingness of respondents to participate in specific activities with training**

| | Count | % |
|---|---|---|
| Evaluate community annotations of a gene family of which you have expert knowledge | 63 | 61.17% |
| Participate in annotating a gene family of interest | 63 | 60.00% |
| Annotate a gene family as a class project | 41 | 41.41% |

**Table 7: Overall bioinformatics skill level**

| | Count | % |
|---|---|---|
| Never used bioinformatics tools | 7 | 6.31% |
| Beginner | 38 | 34.23% |
| Intermediate | 42 | 37.84% |
| Advanced | 24 | 21.62% |

**Table 8: Respondents working with Big Data**

| | Count | % |
|---|---|---|
| Yes, I currently work with Big Data | 85 | 76.58% |
| No, I do not currently work with Big Data | 26 | 23.42% |

**Table 9: Respondents working with Big Data in the next three years**

| | Count | % |
|---|---|---|
| Yes, I will | 100 | 90.09% |
| No, I won't | 11 | 9.91% |

**Table 10: Types of maize research done by survey respondents**

| | Count | % |
|---|---|---|
| Computational and Large-scale Biology | 47 | 43.52% |
| Quantitative Genetics and/or Breeding | 41 | 37.96% |
| Biochemical and Molecular Genetics | 39 | 36.11% |
| Cell and/or Developmental Biology | 36 | 33.33% |
| Population Genetics/Genomics | 30 | 27.78% |
| Transposons and Epigenetics | 16 | 14.81% |
| Cytogenetics | 11 | 10.19% |
| Education and Outreach | 8 | 7.41% |
| Other | 3 | 2.78% |

*n = 108, participants could select more than one option*

**Table 11: Positions held by survey respondents**

| | Count | % |
|---|---|---|
| Researcher (Independent or PI) | 41 | 37.96% |
| Graduate Student | 28 | 25.93% |
| Post-doc | 19 | 17.59% |
| Faculty/Educator | 11 | 10.19% |
| Undergraduate Student | 10 | 9.26% |
| Industry | 5 | 4.63% |
| Other (please specify) | 4 | 3.70% |

*n = 108, participants could select more than one option.*

**S2 Table. Annotation errors and MAKER-P quality metrics for targeted families**

| Gene ID | Gene name | Transcript count | Canonical transcript | Quality - flagged transcript | AED | QI2 | Missing exons | Extra exons | Different exon length/composition | Evidence accounted by other transcripts | Different UTR length |
|---|---|---|---|---|---|---|---|---|---|---|---|
| Zm00001d043179 | PIN9 | 1 | | | 0.01 | 0.5 | | | X | | |
| Zm00001d006082 | PIN11 | 1 | | | 0.12 | 0.3 | X | | | | |
| Zm00001d007395 | ZmGH3-3 | 1 | | | 0.02 | 0.5 | X | | | | |
| Zm00001d011377 | JAR-1 | 3 | T1 | T1 | 0.1 | 0.6 | | | X (T1) | T2 is correct | X (T1/T3) |
| Zm00001d029577 | ZmGH3-1 | 3 | T2 | T2 | 0.06 | 0.5 | X (T1) | | X (T1/T2) | T3 is correct | |
| Zm00001d051514 | ABCB1 | 4 | T2 | T4 | 0.42 | 0.66 | X (T1) | | X (T1/T3) | T2 is correct | X (T1) |
| Zm00001d050259 | ABCA1 | 3 | T2 | T1 | 0.42 | 0.5 | X (T4) | | X (T2) | T3 is correct | |
| Zm00001d041947 | ABCB21 | 1 | | | 0.3 | 0.7 | | | X | | X |
| Zm00001d031871 | ABCB12 | 2 | T1 | T1 | 0.01 | 0.75 | | | X (T1) | T2 is correct | |
| Zm00001d034087 | ORC-5 | 1 | | | 0.22 | 0.6 | X | X | X | | |
| Zm00001d039371 | TCP-33 | 3 | T1 | T1 | 0.12 | 0.5 | | X (T1) | X (T1) | T2 & T3 are correct | |

**S3 Table. Classical genes curated in Apollo**

| Gene ID from Gramene | Error detection method | Transcript count | Canonical transcript | Quality-flagged transcript | AED | QI2 | Missing exons | Extra exons | Non-canonical splice site | Different exon length /composition | Evidence accounted by other transcripts (includes a mix of transcripts) | Different UTR length |
|---|---|---|---|---|---|---|---|---|---|---|---|---|
| Zm00001d026088 | Gene Tree & AED/QI2 | 2 | T2 | T1 | 0.09 | 0.71 | X | | | X | X | |
| Zm00001d003533 | Gene Tree & AED/QI2 | 2 | T1 | T1 | 0.12 | 0.66 | | | X | X | X | |
| Zm00001d045563 | Gene Tree & AED/QI2 | 2 | T1 | T1 | 0.13 | 0.6 | X | | X | X | X | |
| Zm00001d048082 | Gene Tree & AED/QI2 | 4 | T1 | T1 | 0.14 | 0.71 | | | | X | X | X |
| Zm00001d031620 | Gene Tree & AED/QI2 | 2 | T1 | T1 | 0.15 | 0.75 | | X | | X | X | X |
| Zm00001d029074 | Gene Tree & AED/QI2 | 3 | T1 | T1 | 0.17 | 0.66 | | X | X | X | X | |
| Zm00001d008882 | Gene Tree & AED/QI2 | 2 | T1 | T1 | 0.26 | 0.62 | | | | X | | |
| Zm00001d053899 | Gene Tree & AED/QI2 | 4 | T1 | T1 | 0.29 | 0.66 | X | | X | X | X | |
| Zm00001d050265 | Gene Tree & AED/QI2 | 3 | T2 | T3 | 0.29 | 0.75 | | | X | X | | X |
| Zm00001d049995 | Gene Tree & AED/QI2 | 1 | T1 | T1 | 0.29 | 0.75 | | | | X | | X |
| Zm00001d020971 | Gene Tree & AED/QI2 | 2 | T2 | T2 | 0.31 | 0.75 | | X | X | X | | |
| Zm00001d026148 | Gene Tree & AED/QI2 | 2 | T1 | T2 | 0.39 | 0.5 | X | X | | X | | X |
| Zm00001d042445 | AED/QI2 | 1 | T1 | T1 | 0.02 | 0.4 | | | | | | X |
| Zm00001d036242 | AED/QI2 | 1 | T1 | T1 | 0.02 | 0.66 | | | | | | X |

| Gene ID from Gramene | Error detection method | Transcript count | Canonical transcript | Quality-flagged transcript | AED | QI2 | Missing exons | Extra exons | Non-canonical splice site | Different exon length /composition | Evidence accounted by other transcripts (includes a mix of transcripts) | Different UTR length |
|---|---|---|---|---|---|---|---|---|---|---|---|---|
| Zm00001d025355 | AED/QI2 | 1 | T1 | T1 | 0.04 | 0.75 | X | | | | | |
| Zm00001d011277 | AED/QI2 | 1 | T1 | T1 | 0.09 | 0.33 | | | X | X | | |
| Zm00001d019030 | AED/QI2 | 1 | T1 | T1 | 0.13 | 0.5 | X | X | | | | |
| Zm00001d032969 | AED/QI2 | 1 | T1 | T1 | 0.18 | 0.5 | | X | | | | |
| Zm00001d013777 | AED/QI2 | 1 | T1 | T1 | 0.27 | 0.42 | | X | | | | |
| Zm00001d005890 | AED/QI2 | 1 | T1 | T1 | 0.29 | 0.5 | | | | | | X |
| Zm00001d048702 | AED/QI2 | 2 | T1 | T1 | 0.09 | 0.5 | | X | X | X | X | |
| Zm00001d017111 | AED/QI2 | 2 | T1 | T2 | 0.11 | 0.5 | X | | | X | X | |
| Zm00001d018415 | AED/QI2 | 2 | T2 | T1 | 0.11 | 0.66 | X | | | | X | |
| Zm00001d001960 | AED/QI2 | 2 | T1 | T2 | 0.16 | 0.5 | | | | X | | X |
| Zm00001d017288 | AED/QI2 | 2 | T1 | T2 | 0.17 | 0.66 | X | | | X | X | |
| Zm00001d028963 | AED/QI2 | 2 | T1 | T2 | 0.18 | 0.6 | | X | | X | X | |
| Zm00001d034635 | AED/QI2 | 2 | T1 | T2 | 0.19 | 0.66 | X | | | | X | |
| Zm00001d002982 | AED/QI2 | 2 | T1 | T1/T2 | 0.22/ 0.23 | 0.5/ 0.33 | X | | | | X | |
| Zm00001d015366 | AED/QI2 | 2 | T1 | T2 | 0.27 | 0.75 | | | | X | X | |
| Zm00001d003913 | AED/QI2 | 2 | T1 | T1 | 0.28 | 0.75 | X | | | X | | |
| Zm00001d002449 | AED/QI2 | 2 | T1 | T2 | 0.3 | 0.5 | | | | X | X | |
| Zm00001d049610 | AED/QI2 | 2 | T1 | T2 | 0.33 | 0.66 | X | | | X | X | |
| Zm00001d042287 | AED/QI2 | 2 | T1 | T2 | 0.39 | 0.5 | X | | | X | X | |
| Zm00001d021526 | AED/QI2 | 2 | T1 | T2 | 0.39 | 0.66 | | | | X | X | |
| Zm00001d041882 | AED/QI2 | 3 | T2 | T2 | 0.12 | 0.33 | X | X | | | X | |

| Gene ID from Gramene | Error detection method | Transcript count | Canonical transcript | Quality-flagged transcript | AED | QI2 | Missing exons | Extra exons | Non-canonical splice site | Different exon length /composition | Evidence accounted by other transcripts (includes a mix of transcripts) | Different UTR length |
|---|---|---|---|---|---|---|---|---|---|---|---|---|
| Zm00001d024009 | AED/QI2 | 3 | T1 | T2 | 0.12 | 0.7 | | | | X | X | |
| Zm00001d014947 | AED/QI2 | 3 | T1 | T3 | 0.19 | 0.5 | X | X | | X | X | |
| Zm00001d015618 | AED/QI2 | 3 | T1 | T2 | 0.21 | 0.66 | X | | | X | X | |
| Zm00001d023904 | AED/QI2 | 3 | T1 | T1 | 0.24 | 0.75 | | | | X | X | |
| Zm00001d003157 | AED/QI2 | 3 | T1 | T3 | 0.25 | 0.5 | | | | X | X | |
| Zm00001d019565 | AED/QI2 | 3 | T1 | T2/T3 | 0.27/ 0.28 | 0.5 | | | | X | X | |
| Zm00001d012561 | AED/QI2 | 3 | T1 | T3 | 0.3 | 0.5 | X | | | | X | |
| Zm00001d025267 | AED/QI2 | 3 | T2 | T2 | 0.35 | 0.5 | X | | | | X | |
| Zm00001d043175 | AED/QI2 | 3 | T1 | T3 | 0.41 | 0.5 | | | | X | X | |
| Zm00001d049239 | AED/QI2 | 4 | T1 | T2 | 0.29 | 0.5 | | X | | X | X | |
| Zm00001d045735 | AED/QI2 | 4 | T2 | T2 | 0.23 | 0.75 | | X | | | X | |
| Zm00001d050032 | AED/QI2 | 4 | T3 | T2 | 0.25 | 0.72 | X | X | | | | |
| Zm00001d039260 | AED/QI2 | 4 | T2 | T4 | 0.4 | 0.66 | X | | | X | X | |
| Zm00001d014858 | AED/QI2 | 4 | T1 | T2 | 0.37 | 0.66 | X | | | X | X | |
| Zm00001d013631 | AED/QI2 | 4 | T1 | T1 | 0.38 | 0.6 | X | X | | | X | |
| Zm00001d020636 | AED/QI2 | 4 | T1 | T4 | 0.43 | 0.75 | X | | | X | X | |
| Zm00001d050350 | AED/QI2 | 1 | T1 | T1 | 0.17 | 0.75 | X | | | X | | |
| Zm00001d044705 | Gene Triage | 1 | T1 | - | 0.05 | 1 | | | | X | | X |
| Zm00001d052537 | Gene Triage | 1 | T1 | - | 0.05 | 1 | | | | | | X |
| Zm00001d014842 | Gene Triage | 1 | T1 | - | 0.06 | 0 | | X | | | | X |
| Zm00001d020383 | Gene Triage | 1 | T1 | - | 0.06 | 1 | | | | X | | |

| Gene ID from Gramene | Error detection method | Transcript count | Canonical transcript | Quality-flagged transcript | AED | QI2 | Missing exons | Extra exons | Non-canonical splice site | Different exon length /composition | Evidence accounted by other transcripts (includes a mix of transcripts) | Different UTR length |
|---|---|---|---|---|---|---|---|---|---|---|---|---|
| Zm00001d040331 | Gene Triage | 1 | T1 | - | 0.07 | 1 | | | | | | X |
| Zm00001d003006 | Gene Triage | 2 | T1 | - | 0.07 | 1 | | | | | | X |
| Zm00001d036370 | Gene Triage | 1 | T1 | - | 0.08 | 0 | | X | | | | X |
| Zm00001d037737 | Gene Triage | 3 | T1 | - | 0.1 | 1 | X | | | X | X | |
| Zm00001d032922 | Gene Triage | 1 | T1 | - | 0.1 | 1 | | | | | | X |
| Zm00001d045054 | Gene Triage | 1 | T1 | - | 0.11 | 0.8 | X | | | X | | |
| Zm00001d013258 | Gene Triage | 1 | T1 | - | 0.11 | 1 | X | | | X | | X |
| Zm00001d045055 | Gene Triage | 1 | T1 | - | 0.13 | -1 | X | | | X | | |
| Zm00001d037439 | Gene Triage | 1 | T1 | - | 0.13 | 1 | | X | | | | |
| Zm00001d019648 | Gene Triage | 2 | T2 | - | 0.14 | 1 | | | | X | | X |
| Zm00001d042879 | Gene Triage | 4 | T2 | - | 0.16 | 1 | | X | | X | X | X |
| Zm00001d002353 | Gene Triage | 1 | T1 | - | 0.17 | 0 | | | | X | | X |
| Zm00001d047522 | Gene Triage | 4 | T2 | - | 0.17 | 0.8 | X | | | X | X | |
| Zm00001d024698 | Gene Triage | 3 | T2 | - | 0.18 | 1 | X | | | | X | X |
| Zm00001d003804 | Gene Triage | 1 | T1 | - | 0.18 | 1 | | | | | | X |
| Zm00001d017614 | Gene Triage | 4 | T3 | - | 0.21 | 0.77 | | | | X | X | |
| Zm00001d009431 | Gene Triage | 3 | T1 | - | 0.21 | 0.9 | | | | | | X |
| Zm00001d019148 | Gene Triage | 4 | T2 | - | 0.21 | 1 | | | | X | X | |
| Zm00001d043146 | Gene Triage | 2 | T2 | - | 0.24 | 1 | | | | X | | |
| Zm00001d038725 | Gene Triage | 1 | T1 | - | 0.25 | 1 | | | | X | | |
| Zm00001d015450 | Gene Triage | 3 | T1 | - | 0.28 | 1 | X | | | X | X | X |

| Gene ID from Gramene | Error detection method | Transcript count | Canonical transcript | Quality-flagged transcript | AED | QI2 | Missing exons | Extra exons | Non-canonical splice site | Different exon length /composition | Evidence accounted by other transcripts (includes a mix of transcripts) | Different UTR length |
|---|---|---|---|---|---|---|---|---|---|---|---|---|
| Zm00001d041781 | Gene Triage | 4 | T1 | - | 0.28 | 1 | | X | | X | X | |
| Zm00001d051465 | Gene Triage | 3 | T2 | - | 0.3 | 1 | | | | X | X | X |
| Zm00001d006116 | Gene Triage | 1 | T1 | - | 0.31 | 0.83 | X | X | X | X | | |
| Zm00001d051898 | Gene Triage | 1 | T1 | - | 0.35 | 1 | X | | | | X | |
| Zm00001d018535 | Gene Triage | 3 | T3 | - | 0.37 | 1 | | X | | | | |
| Zm00001d032249 | Gene Triage | 2 | T2 | - | 0.51 | 0.6 | | | | X | | X |
| Zm00001d039132 | Gene Triage | 1 | T1 | - | 0.6 | 0 | X | X | X | X | X | X |
| Zm00001d035760 | Gene Triage | 1 | T1 | - | 0.64 | -1 | | | | | | X |
| Zm00001d037386 | Gene Triage | 1 | T1 | - | 0.01 | 1 | | | | X | X | |

**S4 Table. Evaluation of invertases in maize**

| Gene ID | Gene name | Transcript count | Error detection method | Error flagged in trees | Canonical flagged by quality metrics | Main improvements |
|---|---|---|---|---|---|---|
| Zm00001d002830 | INVVR1 | 1 | not flagged | - | no | 9-nt exon (present in V3) |
| Zm00001d014947 | INVVR2 | 3 | AED/QI2 | - | no | - |
| Zm00001d054075 | INVVR3 | 2 | Gene Tree & AED/QI2 | exon loss | yes | extended exons 1-2 (non-canonical junctions) |
| Zm00001d025943 | pseudogene/ novel | 1 | not flagged | - | no | 9-nt exon (OUR FINDING) |
| Zm00001d016708 | INVCW1 | 2 | not flagged | - | no | 9-nt exon (present in V3) |
| Zm00001d003776[a] | INVCW2 | 1 | Gene Tree | 5'-loss | no | 9-nt exon (present in V3); not originally flagged by curators (GT false negative*) |
| Zm00001d025355 | INVCW3 | 1 | AED/QI2 | - | yes | 9-nt exon (present in V3) |
| Zm00001d001941 | INVCW4 | 1 | Gene Tree | exon loss | no | TWO mini-exons: 9-nt & 19-nt (OUR FINDING). Also extended longest exon (wrong in V3) |
| Zm00001d025354 | INVCW5 | 1 | Gene Tree | exon gain | no | 9-nt exon (OUR FINDING) |
| Zm00001d001944 | INVCW6 | 1 | Gene Tree & AED/QI2 | exon gain | yes | 9-nt exon (OUR FINDING) |
| Zm00001d001943 | INVCW7 | 1 | not flagged | - | no | 9-nt exon (present in V3) |
| Zm00001d041991 | INVCW8 | 1 | Gene Tree & AED/QI2 | exon gain | yes | 9-nt exon (present in V3) |

**Legend.**· [a]Annotation errors for **Zm00001d003776** (INVCW2) were not originally detected with the gene tree visualizer. Therefore, the gene is not found in the "improved classical genes", and was included in sensitivity calculations. Omission of the 9-nt exon resulted in this invertase being truncated at its 5'-end.
